# Cortical-brainstem circuitry attenuates physiological stress reactivity

**DOI:** 10.1101/2023.07.19.549781

**Authors:** Sebastian A. Pace, Ema Lukinic, Tyler Wallace, Carlie McCartney, Brent Myers

## Abstract

Exposure to stressful stimuli promotes multi-system biological responses to restore homeostasis. Catecholaminergic neurons in the rostral ventrolateral medulla (RVLM) facilitate sympathetic activity and promote physiological adaptations, including glycemic mobilization and corticosterone release. While it is unclear how brain regions involved in the cognitive appraisal of stress regulate RVLM neural activity, recent studies found that the rodent ventromedial prefrontal cortex (vmPFC) mediates stress appraisal and physiological stress responses. Thus, a vmPFC-RVLM connection could represent a circuit mechanism linking stress appraisal and physiological reactivity. The current study investigated a direct vmPFC-RVLM circuit utilizing genetically-encoded anterograde and retrograde tract tracers. Together, these studies found that stress-reactive vmPFC neurons project to catecholaminergic neurons throughout the ventrolateral medulla in male and female rats. Next, we utilized optogenetic terminal stimulation to evoke vmPFC synaptic glutamate release in the RVLM. Photostimulating the vmPFC-RVLM circuit during restraint stress suppressed glycemic stress responses in males, without altering the female response. However, circuit stimulation decreased corticosterone responses to stress in both sexes. Circuit stimulation did not modulate affective behavior in either sex. Further analysis indicated that circuit stimulation preferentially activated non-catecholaminergic medullary neurons in both sexes. Additionally, vmPFC terminals targeted medullary inhibitory neurons. Thus, both male and female rats have a direct vmPFC projection to the RVLM that reduces endocrine stress responses, likely through the recruitment of local RVLM inhibitory neurons. Ultimately, the excitatory/inhibitory balance of vmPFC synapses in the RVLM may regulate stress reactivity as well as stress-related health outcomes.

**First author profile:** Sebastian Pace is a Ph.D. candidate in the laboratory of Brent Myers at Colorado State University. Before beginning his Ph.D., Sebastian completed his B.S. and M.S. at the University of Texas-El Paso. While completing his M.S. he worked in the Karine Fénelon Lab. Sebastian’s current research project focuses on understanding how corticolimbic inputs to the medullary brainstem regulate sympathetic and neuroendocrine stress responses.

**Key points summary:**

- Glutamatergic efferents from the ventromedial prefrontal cortex target catecholaminergic neurons throughout the ventrolateral medulla.
- Partially segregated, stress-responsive ventromedial prefrontal cortex populations innervate the rostral and caudal ventrolateral medulla.
- Stimulating ventromedial prefrontal cortex synapses in the rostral ventrolateral medulla decreases stress-induced glucocorticoid release in males and females.
- Stimulating ventromedial prefrontal cortex terminals in the rostral ventrolateral medulla preferentially activates non-catecholaminergic neurons.
- Ventromedial prefrontal cortex terminals target medullary inhibitory neurons.

**Graphical Abstract:** 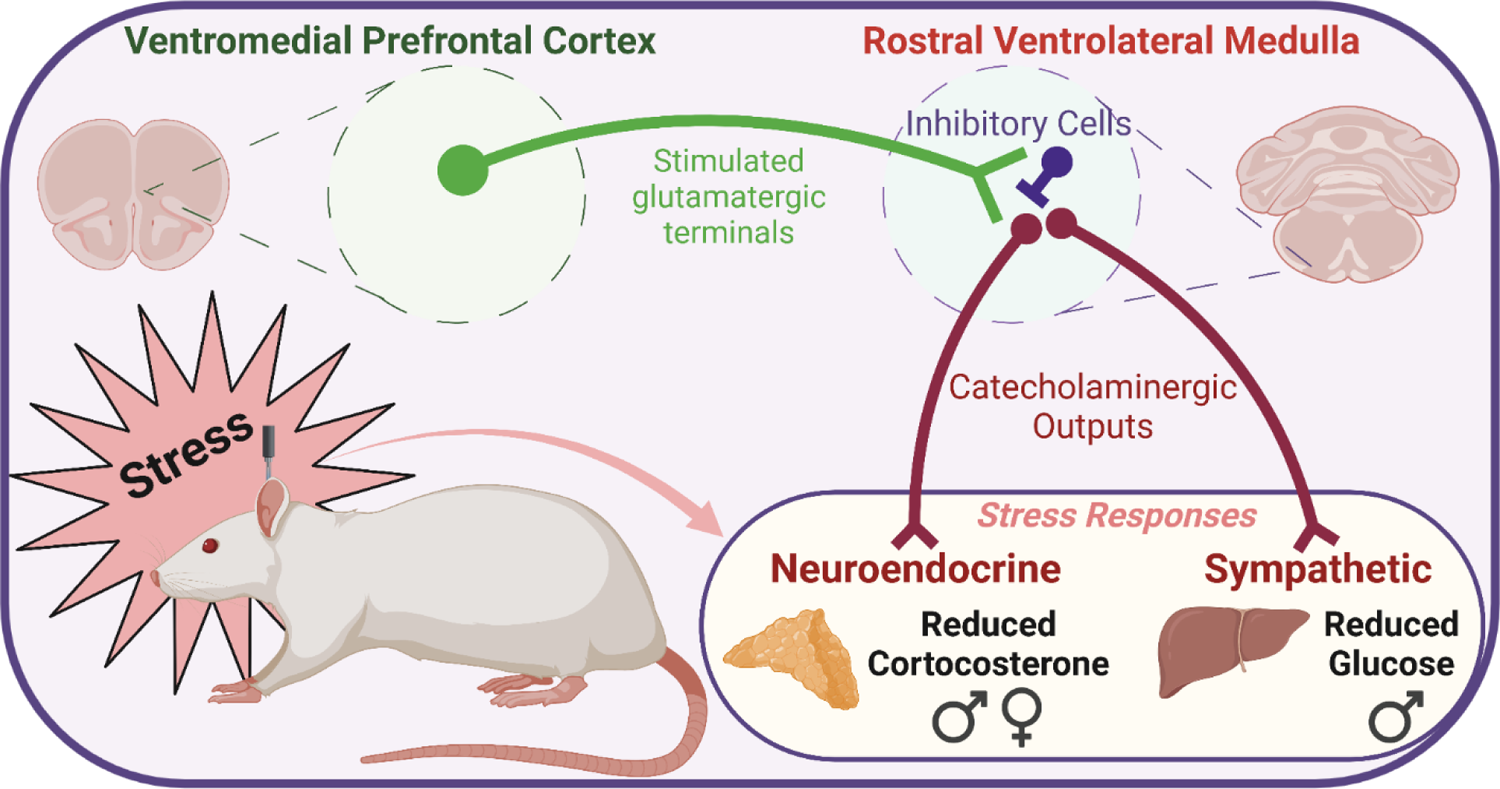

## Introduction

Organismal adaptation to stress is essential for survival. These adaptations are dependent on forebrain structures such as the ventromedial prefrontal cortex (vmPFC) that appraise stressful stimuli and orchestrate appropriate responses (Duncan, 2001; McKlveen et al., 2015; Ulrich-Lai & Herman, 2009). Additionally, stress-related disorders such as depression and post-traumatic stress disorder associate with altered structure and function of the vmPFC (Drevets et al., 1997; Drevets et al., 2008; Hamani et al., 2011; Liotti et al., 2000). In rodents, male vmPFC glutamate neurons promote socio-motivational behaviors and reduce physiological stress responses including hyperglycemia, tachycardia, and corticosterone release (Myers et al., 2017; Schaeuble et al., 2019; Wallace et al., 2021). Notably, vmPFC neurons do not directly project to neurosecretory cells or preganglionic sympathetic neurons, requiring intervening effector(s) to modulate physiological responses (Ulrich-Lai & Herman, 2009). Tract-tracing experiments examined long-range vmPFC projections to the brainstem with some reporting projections to the rostral ventrolateral medulla (RVLM) (Gabbott et al., 2005; Hurley et al., 1991), a key pre-sympathetic nucleus. Moreover, vmPFC neurons innervate catecholaminergic and non-catecholaminergic neurons in the RVLM (Gabbott et al., 2007). However, little is known about the stress responsiveness or function of vmPFC projections to the medulla, nor whether cortical inputs activate the epinephrine/norepinephrine-producing neurons that initiate physiological stress responses.

Brainstem catecholaminergic neurons are evolutionarily-conserved nuclei that facilitate metabolic, neuroendocrine, and autonomic responses to physical and psychological stimuli (Ritter et al., 2019; Stornetta & Guyenet, 2018). Specifically, catecholaminergic neurons in the RVLM activate spinal preganglionic sympathetic neurons to elicit widespread physiological adaptations including vasoconstriction and hyperglycemia (Guyenet et al., 2013). Additionally, ascending RVLM projections target paraventricular hypothalamic neuroendocrine cells that initiate hypothalamic-pituitary-adrenal (HPA) axis glucocorticoid release (Card et al., 2006; Stornetta et al., 2016). Collectively, these physiological effects support a vital homeostatic role for RVLM catecholaminergic neurons (Ritter, 2017). However, the mechanisms regulating RVLM activity during stress have received limited attention. Further, little is known regarding the functional effects of forebrain circuits targeting the RVLM. Here, we examined cortical afferents targeting the RVLM and the neurogenic regulation of stress reactivity by the prefrontal-medullary neural circuit.

To determine if glutamatergic vmPFC projections target RVLM catecholaminergic neurons, an anterograde genetically-encoded vmPFC tract-tracing approach was used to quantify cortical appositions onto epinephrine/norepinephrine-synthesizing neurons throughout the ventrolateral medulla. Further, dual retrograde-transported viruses were used to determine circuit organization and stress responsiveness of cortical-medullary projections. Next, optogenetic stimulation of vmPFC synapses in the RVLM was used to examine motivational behavior as well as glycemic and glucocorticoid responses to stress as end products of the sympathetic and HPA axes, respectively (Bialik et al., 1988; Myers et al., 2014). Additionally, vmPFC-RVLM stimulation tissue was assessed to examine catecholaminergic and non-catecholaminergic cellular activation as well as vmPFC projections onto RVLM inhibitory GABAergic and glycinergic neurons.

## Methods

### Animals

Adult male and female Sprague-Dawley rats (Envigo, Denver, CO) weighing 250-300g and 150-200g, respectively were housed in temperature- and humidity-controlled vivarium with a 12-hour light-dark cycle (lights on at 0600, and off at 1800). Holding rooms were restricted to same-sex conspecific rats. Incoming rats were acclimated to the vivarium for 1-week before the start of the experiment. Water and chow were available ad libitum throughout the experiment. All procedures and protocols were approved by the Institutional Animal Care and Use Committee of Colorado State University (protocol: 1392) and complied with the National Institutes of Health Guidelines for the Care and Use of Laboratory Animals. The experimental procedures used in the current experiments received veterinary consultation and all animals had daily welfare assessments by veterinary and/or animal medical service staff.

### Experimental Design

Experiment 1 was an anterograde tract-tracing experiment comprised of 5 male and 4 female rats for identification of inputs to catecholamine-producing cells. 3 male and female rats from this experiment were also used to for immunolabeling RVLM inhibitory neurons to examine vmPFC terminals appositions. Experiment 2 was a retrograde tract-tracing experiment comprised of 5 male and female rats. Experiment 3 was a vmPFC-RVLM circuit stimulation study that used 2 cohorts of male rats to generate 16 ChR2 and 12 YFP rats. 1 cohort of female rats was used to produce 17 ChR2 and 11 YFP rats. All cohorts were run separately, and all rats underwent real-time place preference (RTPP) and restraint stress. Tissue from Experiment 3 was used for Experiment 4, which examined the neurochemical identity of cells activated by vmPFC-RVLM stimulation.

### Stereotaxic Surgery

Male and female rats were anesthetized with aerosolized isoflurane (1-5%) and administered an analgesic (0.6 mg/kg buprenorphine-SR, subcutaneous) as previously described (Wallace et al., 2021). For anterograde tract-tracing experiments (Experiment 1), rats received bilateral microinjections (males 1.5 µL, females 1 µL) of adeno-associated virus (AAV) constructs (UNC Vector Core, Chapel Hill, NC) in the vmPFC (males: 0.6 mm lateral to midline, 2.7 mm anterior to bregma, and 4.2 mm ventral from dura; females: 0.5 mm lateral to midline, 2.3 mm anterior to bregma, and 4 mm ventral from dura). This experiment used an AAV5-packaged construct to induce expression of yellow fluorescent protein (YFP) under a calcium/calmodulin-dependent protein kinase II alpha (CaMKIIα) promoter. CaMKIIα promoters predominantly induce expression in excitatory glutamatergic neurons (Wood et al., 2019). For retrograde tract-tracing experiments (Experiment 2), rats received 2 separate unilateral microinjections (males 0.2 µL, females 0.15 µL) of retrograde-transported AAV (AAVretro) constructs (Addgene) in the RVLM and CVLM (RVLM-males: +1.9 mm lateral to midline, −12.25 mm posterior to bregma, and −10.4 mm ventral from skull; CVLM-males: +1.85 mm lateral to midline, −12.45 mm posterior to bregma, −12.00 mm ventral from skull, and angled at −8° directed-posteriorly; RVLM-females: +1.8 mm lateral to midline, −12 mm posterior to bregma, and −10.1 mm ventral from skull; CVLM-females: +1.65 mm lateral to midline, −12.40 mm posterior to bregma, −11.3 mm ventral from skull, and angled at −8° directed-posteriorly). An AAVretro construct encoding a mCherry fluorophore targeted the RVLM while an AAVretro encoding a GFP fluorophore targeted the CVLM. Injection laterality was counter-balanced throughout the experiment. AAVretro constructs expressed under a human synapsin promoter (hSyn1). For optogenetic terminal stimulation experiments (Experiment 3), rats received bilateral microinjections (males 1.5 µL, females 1 µL) of AAV constructs (UNC Vector Core) in the vmPFC (males: 0.6 mm lateral to midline, 2.7 mm anterior to bregma, and 4.2 mm ventral from dura; females: 0.5 mm lateral to midline, 2.3 mm anterior to bregma, and 4 mm ventral from dura). AAV5-packaged constructs induced expression of YFP in control rats or channelrhodopsin-2 (ChR2) conjugated to YFP. These viral constructs were under the control of the hSyn promoter as long-range vmPFC projection neurons are overwhelming glutamatergic (Myers et al., 2014). All microinjections used a 25-gauge, 2-µL microsyringe (Hamilton, Reno, NV) and a microinjector (Kopf, Tujunga, CA) at a rate of 5 minutes/µL for vmPFC injections and a rate of 10 minutes/μL for RVLM and CVLM injections. For RVLM injections, the needle was lowered ventrally to −6 mm from the skull and then lowered in −0.5 mm increments every 4 minutes to reduce damage to the respiratory column. The needle was left in place for 10 minutes before and after injections to facilitate viral diffusion. The skin was closed with wound clips that were removed after 2 weeks of recovery. 6 weeks of incubation was given to ensure appropriate viral construct expression (Wallace et al., 2021; Wood et al., 2019).

### Fiber Optic Cannulations

For Experiment 3, male and female rats were anesthetized with aerosolized isoflurane (1-5%) and administered an analgesic (0.6 mg/kg buprenorphine-SR, subcutaneous) and antibiotic (5 mg/kg gentamicin, intramuscular) 4 weeks after microinjections. Unilateral fiber optic cannulas (flat tip, 200 μm diameter, 9.7 mm long for males, 9.4 mm long for females) (Doric Lenses, Québec, Canada) targeted the RVLM (males: +1.82 mm lateral to midline, −12.25 mm posterior to bregma, and −10.2 mm ventral from bregma; females: +1.79 mm lateral to midline, −11.9 mm anterior to bregma, and −9.95 mm ventral from bregma). Fiber optic laterality was counterbalanced, and cannulas were secured via a dental cement headcap (Stoelting, Wood Dale, IL) using dispersed metal screws (Plastics One, Roanoke, VA) as support points. Skin was sutured closed then sutures were removed 10-14 days later. After 2 weeks of recovery, rats underwent 3 days of handling for habituation to tethering.

### Photostimulation Parameters

Light pulses (5.9-6.4 mW, 5 ms pulses, 10 Hz) were delivered through a fiber-optic patch cord (400 µm core diameter, NA = 0.57; Doric Lenses, Québec, Canada) connected to a 473 nm LED driver (Doric Lenses) (Wallace et al., 2021). Optic power was measured using a photodiode sensor (PM160; Thorlabs Inc, Newton, NJ) at the cannula fiber tip in a dark room.

### Estrous Cycle Cytology

Female rats for each experiment were run simultaneously, housed in the same room, and swabbed for estrous cycle cytology. Following experimental assays, vaginal cytology was examined to approximate the estrous cycle stage. A cotton swab dipped in deionized water was used to collect cells from the vaginal canal and roll them onto a glass slide. When dried, slides were viewed under a 10x objective light microscope by a minimum of two blind observers and were categorized as proestrus, estrus, metestrus, or diestrus (Cora et al., 2015; Solomon et al., 2007; Wallace et al., 2021). Any cases with differing estrous stages were resolved by a third blind observer. The distribution of rats in each estrous phase are in Table 1 for each *in vivo* assay.

**Table 1:**
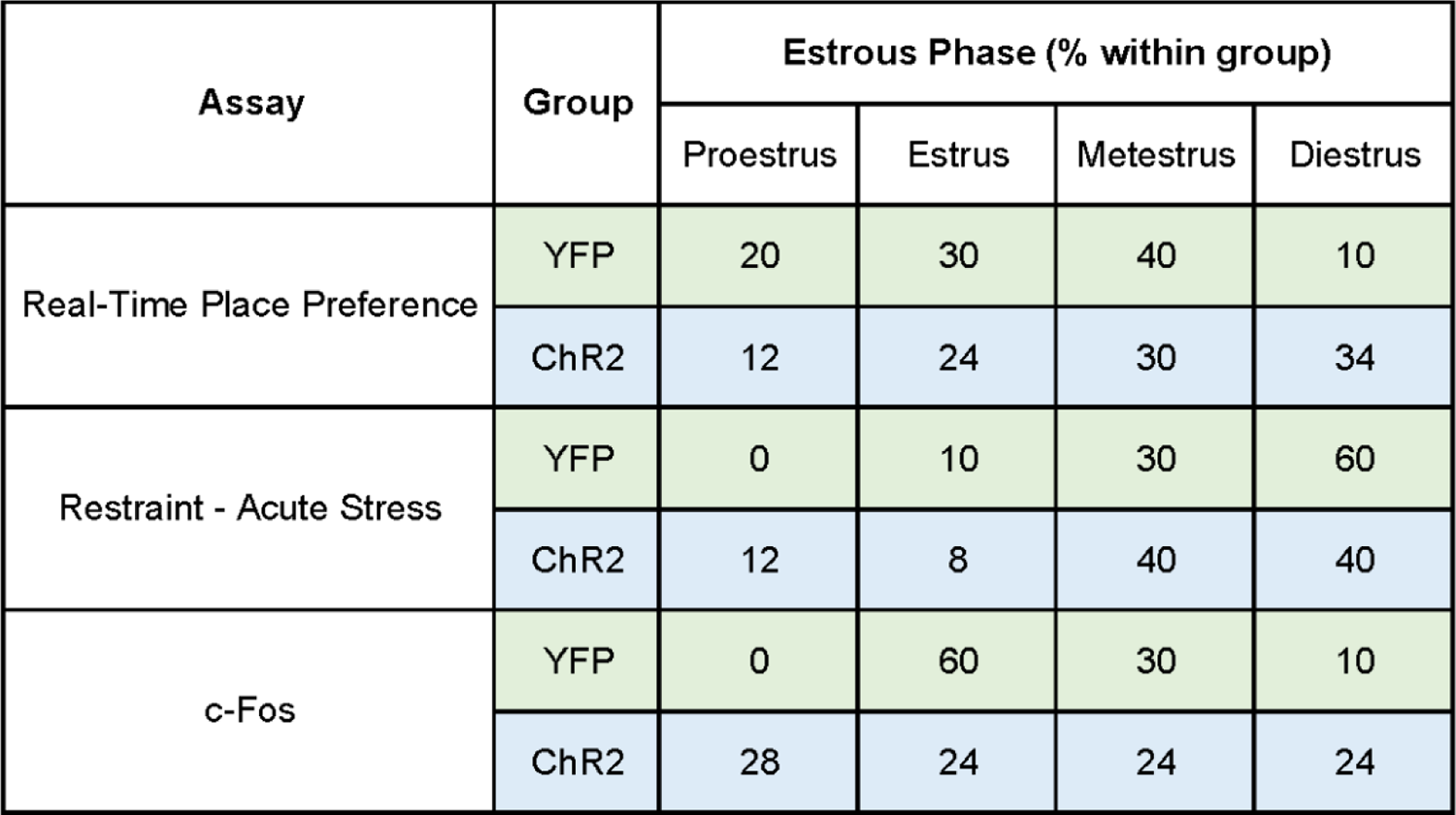
Distribution of estrous cycle phase during testing.

| Assay | Group | Estrous Phase (% within group) |  |  |  |
| --- | --- | --- | --- | --- | --- |
|  |  | Proestrus | Estrus | Metestrus | Diestrus |
| Real-Time Place Preference | YFP | 20 | 30 | 40 | 10 |
|  | ChR2 | 12 | 24 | 30 | 34 |
| Restraint - Acute Stress | YFP | 0 | 10 | 30 | 60 |
|  | ChR2 | 12 | 8 | 40 | 40 |
| c-Fos | YFP | 0 | 60 | 30 | 10 |
|  | ChR2 | 28 | 24 | 24 | 24 |

### Restraint Stress

Restraint stress was used for identifying stress-activated cells (Experiment 2) and to examine neuroendocrine responses to acute stress (Experiment 3). Rats were placed in plastic decapicones (Braintree Scientific, Braintree, MA) and a small window was cut in the plastic to expose the cannula as previously described (Wallace & Myers, 2023). Next, fiber-optic patch cords were attached for optic stimulation throughout the 30-minute restraint. Blood samples were collected via tail clip at the start of restraint with sequential samples taken at 15- and 30-minute timepoints (Myers et al., 2017). At 30 minutes, patch cords were detached and rats returned to the homecage for recovery. Additional blood samples were collected at 60- and 90-minute timepoints. Blood glucose was measured as an indicator of sympathetic outflow to the periphery as acute glucose mobilization is epinephrine-dependent and enhanced by RVLM catecholaminergic stimulation (Zhao et al., 2017). Blood glucose was measured with Contour Next EZ glucometers (Bayer, Parsippany, NJ) and an average of 2 readings were used at each time point. Blood samples were centrifuged at 3000 × g for 15 minutes at 4° C and plasma was stored at −20° C until ELISA analysis. Plasma corticosterone was measured using an ENZO Corticosterone ELISA (ENZO Life Sciences, Farmingdale, NY) with an intra-assay coefficient of variation of 8.4% and an inter-assay coefficient of variation of 8.2% (Bekhbat et al., 2018; Dearing et al., 2021).

### Real-Time Place Preference

The RTPP assay was used to assess valence, or the hedonic quality of vmPFC-RVLM stimulation (Stamatakis & Stuber, 2012). Cannulated rats were attached to fiber-optic patch cord for light delivery and rats were placed in a matte black fiberglass arena (Alpha Plastics and Design, Fort Collins, CO) with two chambers connected by a corridor (chambers: 15” x 15”; corridor: 8” x 6”; entire arena 15” deep). Rats explored the arena for 15 minutes for habituation. During another 15-minute session the next day, rats received 473 nm light pulses when occupying an assigned stimulation chamber. Assigned stimulation chambers were counter-balanced and animal testing was randomized throughout the experiment. Each trial was recorded by a camera mounted above the arena and rat movement was tracked using Ethovision software (Noldus Information Technologies, Leesburg, VA). Tracking software was linked to the LED drivers to automate optics during the assay by a mini USB-IO box (Noldus Information Technologies). The time rats spent on the stimulation side was divided by the total time and multiplied by 100 to calculate the percentage of time spent on the stimulation side.

### Tissue Collection

After experiments, all rodents were anesthetized using sodium pentobarbital (≥100 mg/kg, intraperitoneal) and then transcardially perfused with 0.9% saline followed by 4% phosphate-buffered paraformaldehyde. Brains were post-fixed in paraformaldehyde overnight and then stored in 30% sucrose at 4°C. Brains were subsequently sectioned (30 μm thick 1:12 serial coronal sections) and stored in cryoprotectant solution at −20 °C until immunohistochemistry. Rats in experiment 2 were exposed to a 30 min restraint stressor 90 min prior to euthanasia. Rats involved in experiment 3 received optical stimulation before tissue collection. Rats were tethered to a fiber optic patch cord and received 5 min of optic stimulation (5.9-6.4 mW, 5 ms pulses, 10 Hz) followed by 90 min of recovery for immediate-early gene (c-Fos) expression before euthanasia, as described above.

### Immunohistochemistry

For fluorescent labeling of dopamine beta-hydroxylase (DBH), coronal sections were removed from the cryoprotectant and rinsed in phosphate buffered saline (PBS) (5 x 5 min) at room temperature. Sections were then moved to blocking solution (PBS, 0.1% bovine serum albumin, 0.2% Triton X-100) for 1 hr. Sections were then incubated overnight in mouse anti-DBH primary antibody (1:2500 in blocking solution, MAB394, RRID: AB_94983; MilliporeSigma, Burlington, MA). Next, sections were rinsed in PBS (5 x 5 min) and then incubated in donkey anti-mouse Cy3 secondary antibody (1:200 in PBS, 715-165-020, RRID: AB_2340811; Jackson ImmunoResearch, West Grove, PA) for 1 hr. The tissue was then washed in PBS (5 x 5 min) and placed into a DAPI stain solution (300 nM in PBS, D3571, RRID: AB_2307445; ThermoFisher Scientific, Portsmouth, NH) for 10 minutes. After another PBS wash (5 x 5 min), the tissue was mounted in polyvinyl medium and cover-slipped for imaging.

For fluorescent labeling of c-Fos (Experiment 3), coronal sections were rinsed in PBS (5 x 5 min) and moved to blocking solution (PBS, 0.1% bovine serum albumin, 0.2% Triton X-100) for 1 hr. Sections were incubated for 48 hours in rabbit anti-c-Fos primary antibody (1:1000 in blocking solution, 226_003, waiting on RRID approval; Synaptic Systems, Goettingen, Germany). Next, sections were rinsed in PBS (5 x 5 min) and incubated in donkey anti-rabbit Cy5 secondary antibody (1:1000 in PBS, 711-175-152, RRID: AB_2340607; Jackson ImmunoResearch) for 1 hr. After, the tissue was washed in PBS (5 x 5 min), mounted in polyvinyl medium, and cover-slipped for imaging. For retrograde tract-tracing experiments (Experiment 2), a different version of the c-Fos antibody (226_008, Synaptic Solutions, Goettingen, Germany) was used due to the original version being discontinued.

When double labeling DBH and c-Fos (Experiment 4), sections were rinsed in PBS (5 x 5 min) then moved to blocking solution (PBS, 0.1% bovine serum albumin, 0.2% Triton X-100) for 1 hr. Subsequently, sections were incubated overnight in mouse anti-DBH primary antibody. Next, sections were rinsed in PBS (5 x 5 min) then incubated in donkey anti-mouse Cy3 secondary antibody for 1 hr. The tissue was then washed in PBS (5 x 5 min) and blocking solution (PBS, 0.1% bovine serum albumin, 0.2% Triton X-100) for 1 hr. Sections were incubated for 48 hours in rabbit anti-c-Fos primary antibody. Next, sections were rinsed in PBS (5 x 5 min) and incubated in donkey anti-rabbit Cy5 secondary antibody for 1 hr. Lastly, the tissue was washed in PBS (5 x 5 min), mounted in polyvinyl medium, and cover-slipped for imaging.

To visualize GABA, sections were retrieved, rinsed, and incubated for 4 hr in blocking solution (7.5% normal goat serum, 6% BSA, 1.5% normal donkey serum in 50 mM KPBS). Sections were then incubated in rabbit anti-GABA primary antibody (1:250 in blocking solution, AB141, RRID: AB_11214017; MilliporeSigma) for 60 h. Primary antibody labeling was amplified with biotinylated goat anti-rabbit IgG for 2 hr (1:500 in PBS, BA-1000, RRID: AB_2313606; Vector Laboratories; Burlingame, CA) followed by Vectastain ABC Solution for 1 hr (1:500 in PBS, PK-7100, RRID: AB_2336827; Vector Laboratories) then Cy3-conjugated streptavidin for 1 hr (1:500 in PBS, 016-160-084, RRID: AB_2337244; Jackson ImmunoResearch). Finally, the tissue was washed in PBS (5 x 5 min), mounted in polyvinyl medium, and cover-slipped for imaging.

For immunolabeling glycine transporter 2 (GlyT2), sections were rinsed and blocked for 1 hr (PBS, 0.1% bovine serum albumin, 0.2% Triton X-100). Sections were incubated for 48 hours in rabbit anti-GlyT2 primary antibody (1:1000 in blocking solution, Af1290, RRID: AB_2571606; Frontier Institute, Japan, Shinkonishi). Sections were then rinsed in PBS (5 x 5 min) and incubated in donkey anti-rabbit Cy5 secondary antibody (1:1000 in PBS, 711-175-152, RRID: AB_2340607; Jackson ImmunoResearch) for 1 hr. After, the tissue was washed in PBS (5 x 5 min), mounted in polyvinyl medium, and cover-slipped for imaging.

### Microscopy

All fluorescent microscopy used a Zeiss Axio Imager Z2 microscope (Carl Zeiss Microscopy, Jena, Germany) and the corresponding ZEN 2.6 blue edition software (Carl Zeiss Microscopy). To determine injection placement, YFP was imaged using the 10x objective, while YFP and DBH dual fluorescence imaging used a 63x objective and 0.5-μm thick optical sectioning to produce Z-stacks. For mapping AAVretro-mCherry and -GFP injections and labeled cells, 20x tiled images were taken. Co-localization was defined as purple or yellow fluorescence from the overlap between labeled mCherry or GFP terminals and c-Fos Cy5. RVLM cannula placements were mapped using a 10x tiled slide scan. For c-Fos and c-Fos/DBH quantification after stimulation, 10x tiled images were acquired to determine if nuclear c-Fos labeling was surrounded by the cytosolic DBH labeling. Lastly, GABA and GlyT2 were imaged with a 63x objective and 0.5-μm thick optical sectioning. In all imaging cases, an off-channel filter was used to exclude auto-fluorescent cells that may affect results.

### Image Analysis

For anterograde tract-tracing experiments (Experiment 1), Carl Zeiss Images (CZIs) were imported to a computer equipped with Imaris 8.1.2 (Oxford Instruments, Oxford, UK) to identify and quantify DBH-labeled neurons and YFP-expressing vmPFC terminals. Further, high-magnification 3-D imaging enabled identification of putative terminal appositions on medullary cell bodies, or YFP-expressing fibers overlapping with DBH-labeled neurons. For retrograde tract-tracing experiments (Experiment 2), tiled CZI images were analyzed using ImageJ Fiji (ver. 1.51N) to quantify RVLM- and CVLM-projecting vmPFC neurons. For c-Fos quantification of vmPFC-RVLM terminal stimulation cases (Experiment 3), ImageJ Fiji was utilized to quantify c-Fos-labeled cells in the RVLM. For c-Fos/DBH colocalization experiments (Experiment 4), ImageJ Fiji was used to quantify the number of cells expressing c-Fos and DBH separately or together. Colocalization was defined by nuclear c-Fos-Cy5 signal surrounded by DBH-Cy3 signal.

### Neuroanatomy

To anatomically delineate mPFC, the bregma location and area delineations of each tissue section were defined according to the Brain Maps III: Structure of the Rat Brain (Swanson, 2004). The atlas was used to identify the anterior forceps of the corpus callosum as the lateral boundary and the coronal midline as the medial boundary. The rostral-caudal emergence of the corpus callosum was used to divide the subregions from dorsal to ventral and the subependymal zone guided the identification of the ventral vmPFC boundary. To delineate the RVLM, The Rat Brain atlas (Paxinos & Watson, 2006) was used for area delineations and landmark identification throughout the brainstem. The RVLM lacks distinct cytoarchitecture, therefore, DBH-labeling was repeatedly used to identify the RVLM while landmarks such as the facial nucleus served as the rostral boundary, spinal trigeminal nucleus served as the lateral boundary, and lateral portions of the inferior olive guided distinguishing the medial boundary. Regarding VLM subregions, The Rat Brain atlas (Paxinos and Watson, 2006) defines the whole VLM as −12.00 to −15.00 mm from bregma, with catecholaminergic populations transitioning from C1, C1/A1, and A1 in a rostro-caudal orientation. We used these catecholaminergic populations to define what we considered RVLM (−12.00 to −13.56 mm posterior to bregma), intermediate VLM (−13.68 to −14.16 mm posterior to bregma), and CVLM (Bregma −14.28 to −15.00 mm posterior to bregma) (Li et al., 2018). Notably, our classification regards the RVLM as containing catecholaminergic neurons that have bulbospinal as well as ascending projections. Further, templates of rat brain coronal sections from Brain Maps III (Swanson, 2004) were used to illustrate virus and cannula placement.

### Data Analysis

Data are expressed as mean ± standard error of the mean. Data were analyzed using Prism 9 (GraphPad, San Diego, CA), with statistical significance set at p < 0.05 for all tests. A one-way ANOVA was used to analyze the number of cells with appositions and the number of appositions per cell across VLM subregions. For retrograde tract-tracing experiments, a one-way ANOVA was used to analyze density of medullary-projecting vmPFC neurons. Stimulation induced c-Fos counts were analyzed with Welch’s unpaired t-test comparing treatment groups. RTPP stimulation preference was assessed via Welch’s unpaired t-test comparing treatment groups. Corticosterone and glucose measured during restraint stress were analyzed using a repeated mixed-effects analysis with treatment and time as factors. If significant main or interaction effects are present, then a Fisher’s post-hoc test was used. DBH/c-Fos colocalization analyses used a Welch’s unpaired t-test. Throughout all experiments, comparisons were limited to within-sex as differences in construct expression across sexes confound comparisons.

## Results

### vmPFC Glutamate Projections to Catecholaminergic VLM Neurons

Microinjections of the viral construct expressing YFP under the CaMKIIα promoter targeted the vmPFC of adult male and female rats (Fig.1A) and placement was mapped using a rat brain atlas (Fig. 1C, G). Anterograde injections typically infected fewer vmPFC starter neurons in females compared to males (average surface area of YFP injection site: males 2.149 mm^2^; females, 1.397 mm^2^). Following viral transduction, YFP-expressing vmPFC fibers were observed in the ventrolateral medulla (VLM) (Fig. 1B). In the VLM, neurons were immunolabeled for the epinephrine- and norepinephrine-synthesis enzyme, DBH. Further, given the lack of cytoarchitectonic parcellations or laminations in the VLM, the presence of DBH+ neurons and the use of regional landmarks (facial, trigeminal, and olivary nuclei) helped to define VLM spatial boundaries. Next, putative appositions of YFP-expressing vmPFC fibers on catecholaminergic VLM cell bodies were quantified throughout the VLM in male (Fig. 1D) and female (Fig. 1H) rats. In males, vmPFC fibers apposed the majority of DBH+ VLM neurons in the rostral, intermediate, and caudal portions of the VLM (Fig. 1E). The quantity and density of vmPFC appositions on catecholaminergic neurons did not vary by VLM subregion, (Fig. 1E, F) [quantity of appositions, n = 6, one-way ANOVA (F(2,12) = 0.43, p = 0.66); number of appositions, n = 5, one-way ANOVA (F(2,12) = 0.57, p = 0.58). Similarly, in females, YFP-expressing vmPFC fibers apposing DBH+ neurons were seen across the VLM (Fig. 1H, I) [F(2,6) = 0.22, p = 0.98]. Again, appositions on DBH+ neurons were not significantly different by VLM subregion (Fig. 1J) [F(2,6) = 0.052, p = 0.95]. In all, these anterograde tract-tracing studies revealed glutamatergic vmPFC neurons target catecholaminergic neurons throughout the VLM in male and female rats.

**Figure 1.**
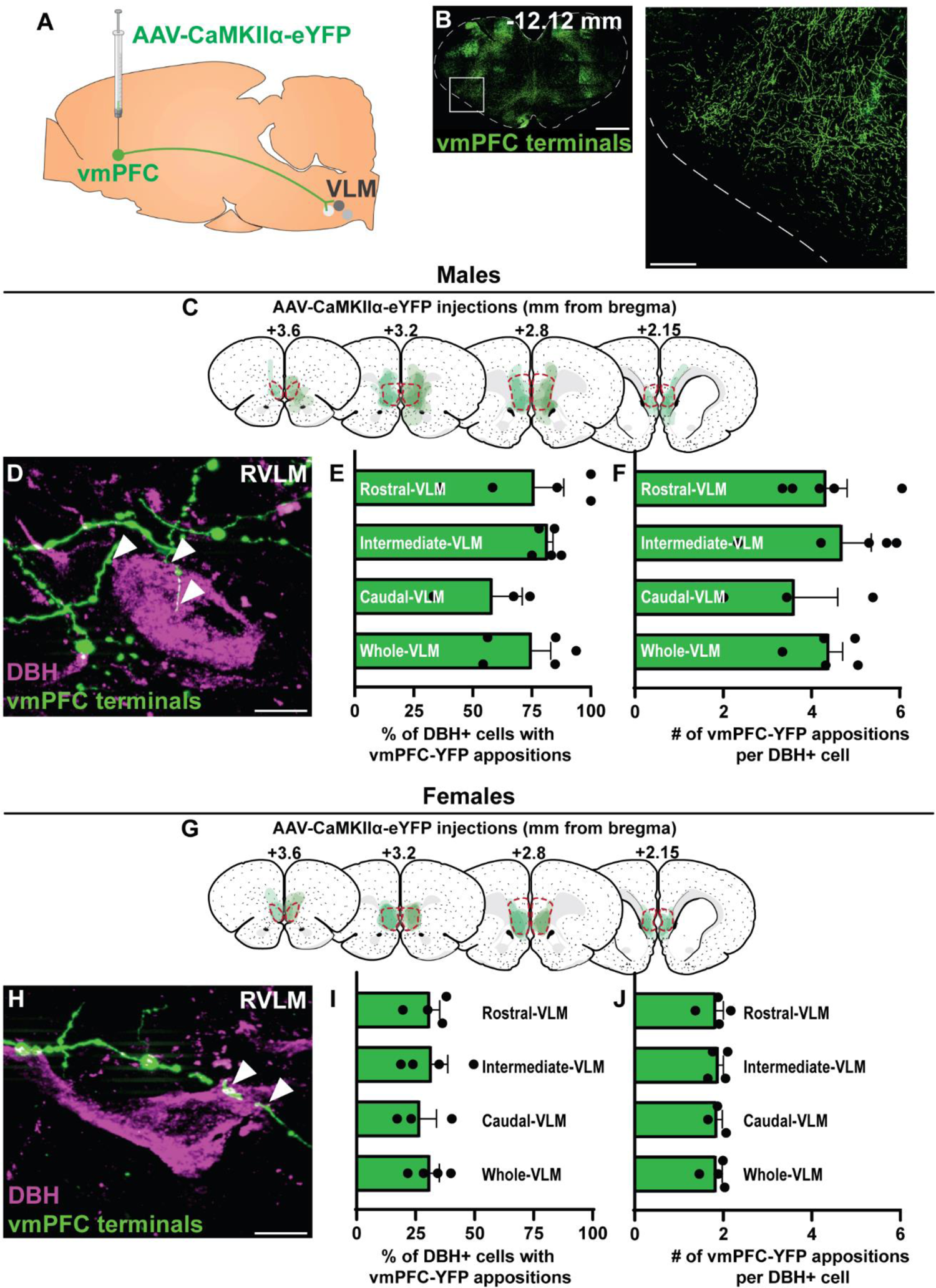
vmPFC neurons project to catecholaminergic neurons throughout the VLM. AAV-YFP was injected into the vmPFC and terminal expression of YFP was identified in the VLM, scale bar: 1 mm, inset scale bar: 250 μm (A, B). Male rat microinjections were mapped onto Swanson Rat Brain Atlas (3rd edition) coronal sections with the vmPFC outlined and rostral-caudal distance to bregma noted above sections (C). YFP-expressing terminals apposed catecholaminergic neurons, labeled with DBH, arrowheads denote appositions, scale bar: 5 μm (D). vmPFC terminals apposed the majority of male DBH+ neurons throughout the VLM subregions (E). The number of vmPFC terminal appositions per each DBH+ VLM neuron was similar throughout the VLM (F). Female rat microinjections were mapped onto coronal sections with the vmPFC (red outline) and rostral-caudal distance to bregma noted above sections (G). Putative appositions of YFP-expressing on DBH+ neurons were observed, arrowheads denote appositions, scale bar: 5 μm (H). vmPFC terminal appositions similarly targeted DBH+ neurons throughout the female VLM and individual subregions (I). The number of vmPFC terminal appositions per each DBH+ VLM neuron was similar throughout the VLM (J). AAV-CaMKIIα-eYFP: adeno-associated virus to express YFP under a calcium/calmodulin-dependent protein kinase II alpha promoter, DBH: dopamine beta-hydroxylase, RVLM: rostral ventrolateral medulla, vmPFC: ventromedial prefrontal cortex, YFP: yellow fluorescent protein.

### Organization and Stress Responsiveness of RVLM- and CVLM-Projecting vmPFC Neurons

The parallel, divergent, or mixed circuit organization of stress-activated vmPFC ensembles that target the RVLM and CVLM was investigated. Dual injections of retrograde-transported viruses separately targeted the RVLM and CVLM in the same subjects (Fig. 2A). AAVretro-mCherry was injected into the RVLM and AAVretro-GFP was injected into the CVLM (Fig. 2B). Viral injection spread and placement were mapped using a rat brain atlas (Fig. 2D, G). Here, the male and female vmPFC was surveyed for RVLM- and CVLM-projecting neurons, as well as co-labeling of c-Fos in response to stress (Fig. 2C). In males, no differences were seen in the density of vmPFC cells projecting to the RVLM and CVLM individually, as well as both the RVLM and CVLM [n = 3/group, one-way ANOVA: F(2,6) = 0.33, p = 0.73] (Fig. 2E). Further, the percentage of stress-reactive vmPFC neurons did not change across neuroanatomical targets [n = 3/group, one-way ANOVA: F(2,6) = 0.14, p = 0.87] (Fig. 2F). Similar trends were seen in females. No differences were evident between the density of vmPFC cells projecting to differing medullary areas [n = 3/group, one-way ANOVA: F(2,9) = 1.07, p = 0.38] (Fig. 2H). Additionally there were no differences between the percentage of stress-reactive vmPFC neurons across neuroanatomical targets [n = 3/group, one-way ANOVA: F(2,9) = 0.33, p = 0.73] (Fig. 2I). In all, stress-reactive vmPFC cells targeted the RVLM and CVLM through both parallel and divergent pathways.

**Figure 2.**
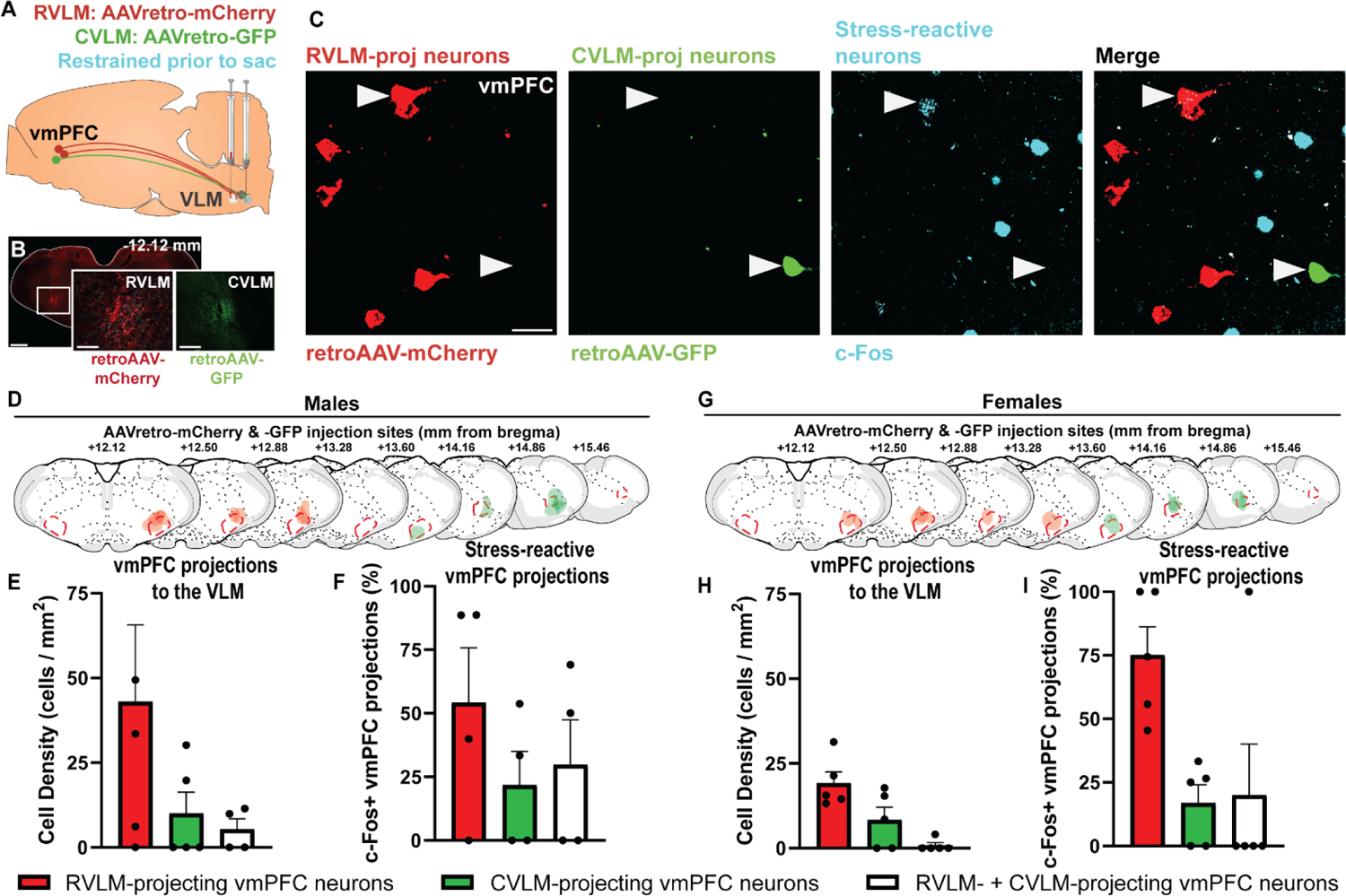
Stress-activated vmPFC neurons target the RVLM and CVLM. AAVretro constructs expressing mCherry or GFP were injected into the RVLM and the CVLM, respectively, scale bar: 1 mm, inset scale bar: 250 μm (A, B). mCherry- and GFP-labeled neurons were present in the vmPFC, as well as c-Fos+ cells following restraint stress, scale bar: 10 μm (C). Male and female rodent microinjections were mapped onto Swanson Rat Brain Atlas (3rd edition) coronal sections with the RVLM and CVLM outlined (D, G). In male rats, the density of vmPFC neurons projecting to the RVLM, CVLM, and both regions simultaneously were not statistically different (E). VLM-projecting neurons were stress-reactive across the vmPFC (F). In female rats, the density of vmPFC neurons projecting to the RVLM, CVLM, and both regions simultaneously were not significantly different (H). Female VLM-projecting neurons throughout the vmPFC were stress-reactive (I). AAVretro: retrograde-traveling adeno-associated virus, CVLM: caudal ventrolateral medulla, GFP: green fluorescent protein, RVLM: rostral ventrolateral medulla, vmPFC: ventromedial prefrontal cortex.

### Optogenetic vmPFC-RVLM Circuit Stimulation

To determine the functional effects of the vmPFC-RVLM circuit, we used an optogenetic terminal stimulation strategy (Fig. 3A-D). Stimulation increased RVLM c-Fos+ cell density (# of c-Fos+ cells /mm^2^) of ChR2 compared to YFP controls, in both male and female rats [males (n = 4-5/group, unpaired t-test: ChR2 vs YFP t(7) = 3.70, p = 0.0076); females (n = 4-6/group, unpaired t-test: ChR2 vs YFP t(7) = 4.02, p = 0.0051)] (Fig. 3E-J).

**Figure 3.**
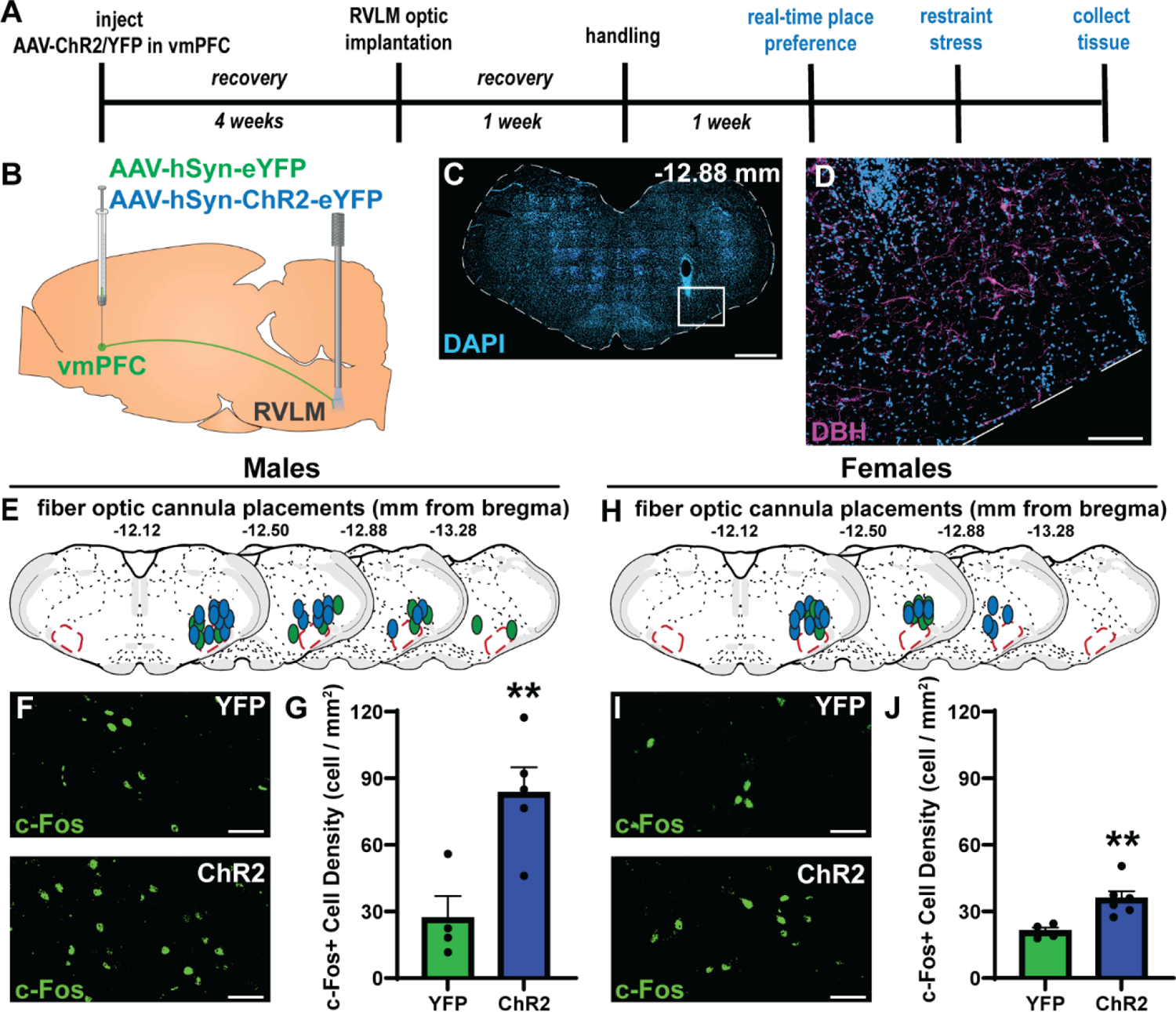
vmPFC terminals stimulated in RVLM increase c-Fos+ neurons. vmPFC-RVLM terminal stimulation experiment timeline (A). AAV-YFP or -ChR2 was injected into the vmPFC and a light-emitting optic fiber was implanted dorsal to the RVLM to stimulate vmPFC synapses at the vmPFC, scale bar: 1 mm, inset scale bar: 200 μm (B-D). Male and female counterbalanced unilateral cannulations were mapped onto Swanson Rat Brain Atlas (3rd edition) coronal sections with the RVLM outlined (E, H). vmPFC-RVLM stimulation increased immunoreactive c-Fos cells in the RVLM of male and female rats, scale bars: 50 μm (F, G, I, J). AAV-hSyn-eYFP: adeno-associated virus to express YFP under a synapsin promoter, AAV-hSyn-ChR2: adeno-associated viral package to express ChR2 under the under a synapsin promoter, ChR2: channelrhodopsin-2, DBH: dopamine beta-hydroxylase, RVLM: rostral ventrolateral medulla, vmPFC: ventromedial prefrontal cortex, YFP: yellow fluorescent protein. * p<0.05, ** p<0.01.

### Motivational Valence of vmPFC-RVLM circuit Stimulation

We examined whether vmPFC-RVLM circuit stimulation induces preference or avoidance behavior using the RTPP assay (Fig. 4A). Here, the ChR2 and YFP groups spent the similar amounts of time in both chambers of the RTPP arena [males (n = 11-17/group, unpaired t-test: ChR2 vs YFP t(30) = 0.26, p = 0.79); females (n = 12-16/group, unpaired t-test: ChR2 vs YFP t(26) = 0.78, p = 0.44)] (Fig. 4B, C). Further, no locomotive phenotype was observed in male or female rats [males distance (n = 11-17/group, unpaired t-test: ChR2 vs YFP t(26) = 0.48, p = 0.63); females distance (n = 12-16/group, unpaired t-test: ChR2 vs YFP t(26) = 0.25, p = 0.80)] (Fig. 4D, E). Thus, although vmPFC mediates affective and reward behaviors (Fuchikami et al., 2015; Pace et al., 2020; Wallace et al., 2021), the vmPFC-RVLM circuit has neutral valence.

**Figure 4.**
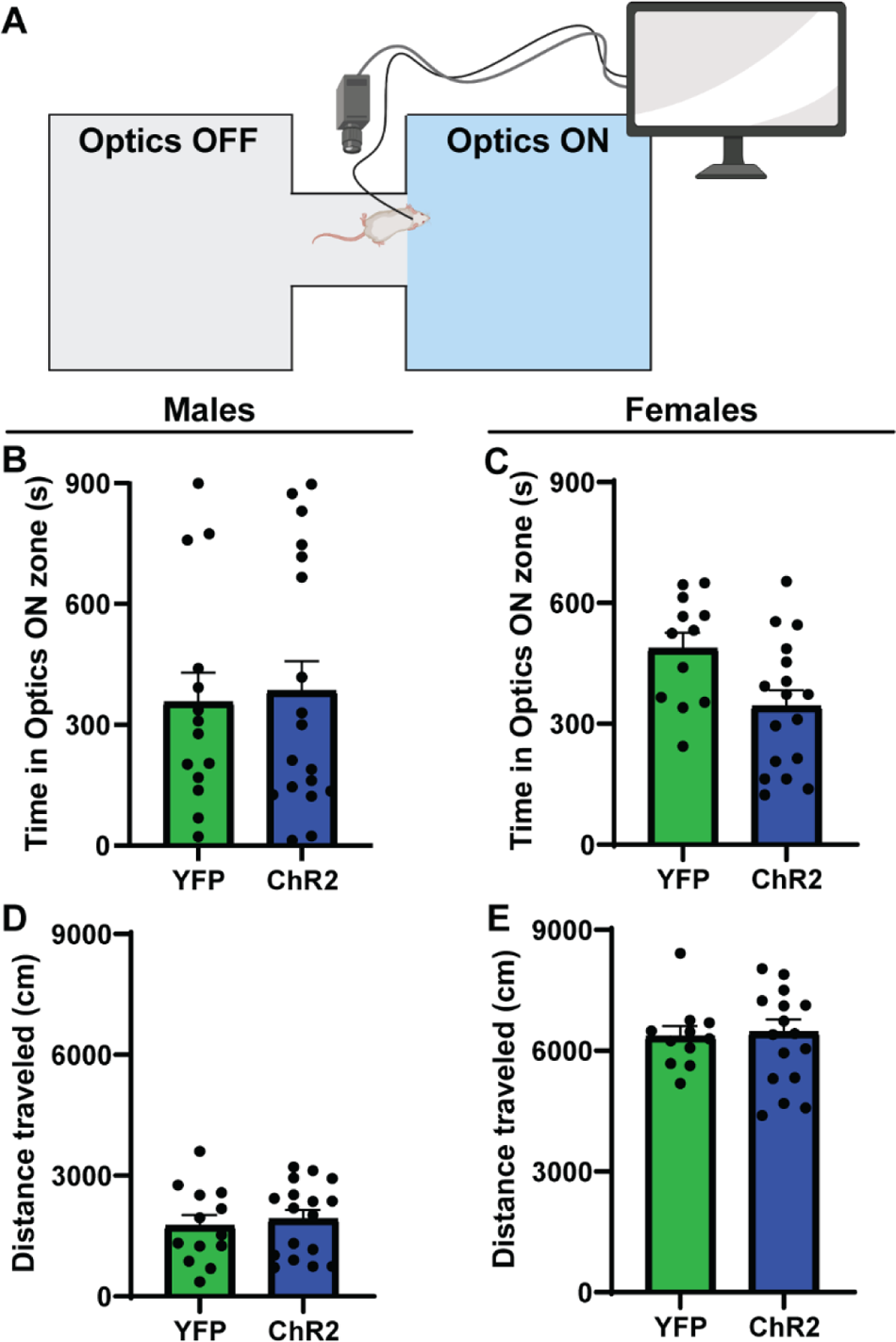
Stimulating the vmPFC-RVLM circuit does not change preference behavior. Cannulated rats underwent a real-time place preference assay in a 2-chamber arena with a connecting space (A). Male and female rats did not show preference or aversion for vmPFC-RVLM stimulation (B, C). Male and female ChR2 groups had no differences in total locomotion relative to YFP controls (D, E).

### vmPFC-RVLM Circuit Regulation of Stress Reactivity

Next, we sought to examine physiological regulation by the vmPFC-RVLM pathway by activating the circuit during stress. We assessed sympathetic and neuroendocrine stress responses by measuring blood glucose and plasma corticosterone, respectively. In males, glucose levels were decreased in ChR2 rats compared to controls at the 30 minute time point of restraint stress [n = 11–17/group, mixed-effects: time F(4,125) = 17.51, p < 0.0001; time x ChR2 F(4,125) = 2.79, p = 0.029; 30 min ChR2, p = 0.0088] (Fig. 5A). Additionally, corticosterone levels were decreased in the ChR2 group following stress (Fig. 5B) at the 60-minute time point [n = 11–17/group, mixed-effects: time F(4,80) = 10.26, p < 0.0001; 60 min ChR2, p = 0.035].

**Figure 5.**
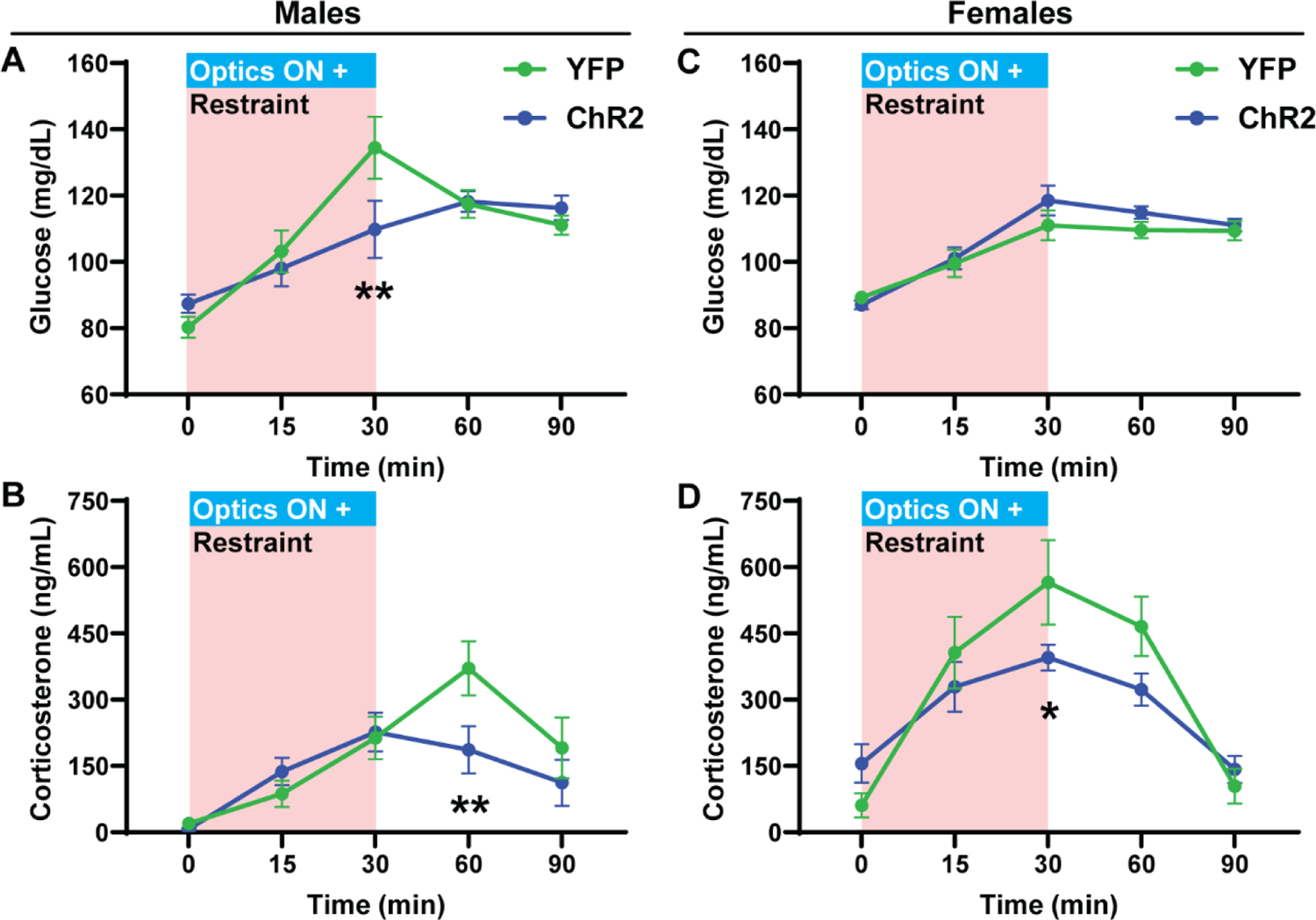
Stimulating the vmPFC-RVLM circuit during restraint stress blunts corticosterone release in both sexes and glucose mobilization in males. Male ChR2 rats had blunted stress-induced glucose mobilization (A). Similarly, male ChR2 rats had reduced corticosterone stress responses (B). Female ChR2 and YFP groups had similar glycemic responses to stress (C). Female ChR2 rats had reduced stress-induced corticosterone secretion (D). * p<0.05, ** p<0.01.

**Figure 6.**
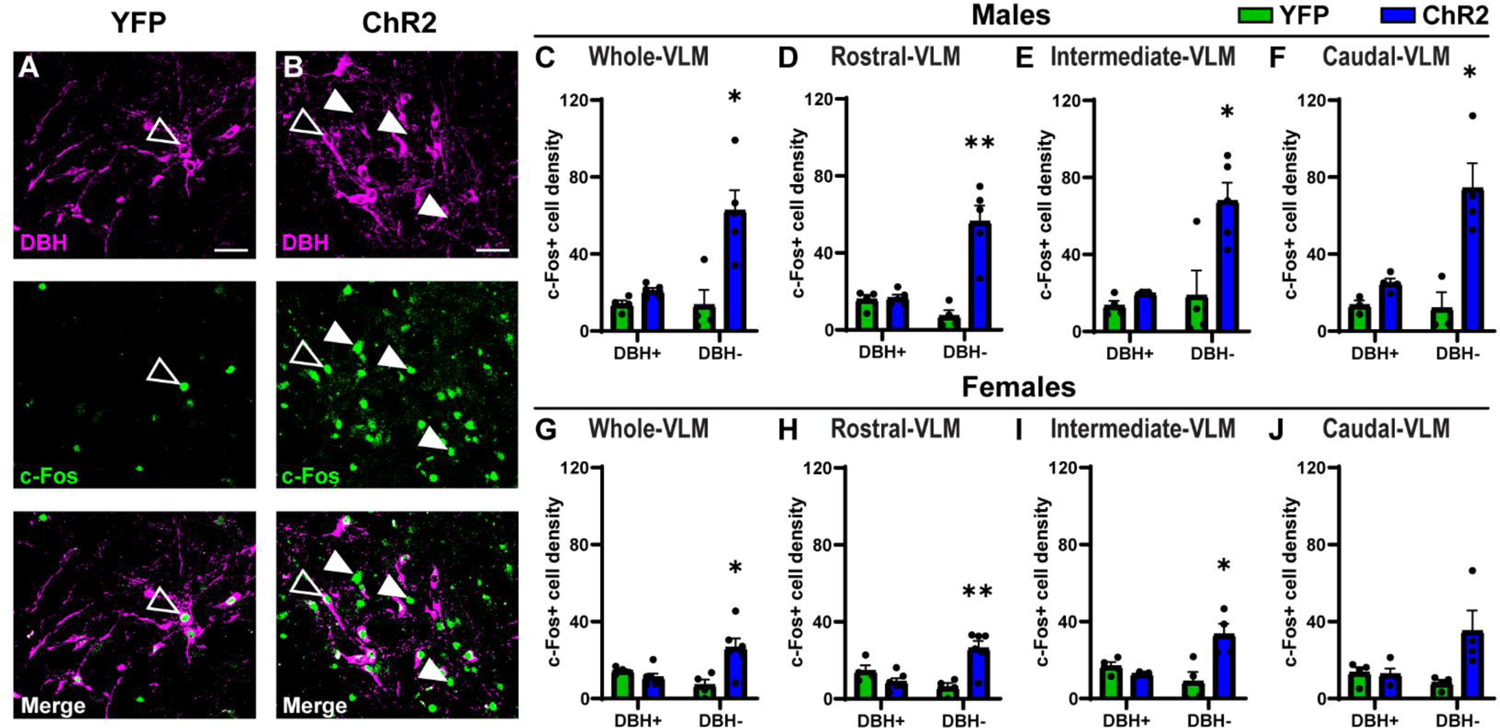
Stimulating the vmPFC-RVLM circuit preferentially activates non-catecholaminergic neurons in the ventrolateral medulla. YFP and ChR2 groups were immunolabeled for DBH and c-Fos, DBH+ and c-Fos+ cells are marked by unfilled arrowheads, DBH- and c-Fos+ cells marked by filled arrowheads, scale bars: 50 μm (A, B). Male ChR2 rats expressed increased c-Fos in DBH- neurons throughout the VLM (C-F). Female ChR2 rats expressed increased c-Fos in DBH- neurons throughout the VLM (G-J). * p<0.05, ** p<0.01.

In contrast, glucose levels were not different at any timepoint (Fig. 5C) in female rats [n = 12–16/group, mixed-effects: time F(4,121) = 19.71, p > 0.05]. However, corticosterone was significantly decreased (Fig. 5D) at the 30-minute time point in female ChR2 rats compared to YFP controls [n = 12–16/group, mixed-effects: time F(4,116) = 19.71, p < 0.0001; 30 min ChR2, p = 0.025]. These results indicate that activation of the vmPFC-RVLM pathway reduced stress-induced glucocorticoid release in both sexes and stress-induced hyperglycemia in males.

### The vmPFC-RVLM Circuit Preferentially Activates Non-Catecholaminergic Cells

Tissue was collected after vmPFC-RVLM stimulation and immunolabeled for DBH and c-Fos to determine if circuit stimulation activated catecholaminergic RVLM neurons (Fig. 5A, B). In males, c-Fos+ cell density (# of c-Fos+ cells / mm^2^) in DBH+ neurons was comparable in YFP and ChR2 groups [whole VLM (n = 4-5/group, unpaired t-test: ChR2 vs YFP t(7) = 2.69, p = 0.07] (Fig. 5C). Further, VLM DBH+ cells had no differences in c-Fos expression within VLM subregions [rostral VLM (unpaired t-test: ChR2 vs YFP t(6) = 0.28, p > 0.99); intermediate VLM (unpaired t-test: ChR2 vs YFP t(3) = 2.48, p = 0.17); caudal VLM (unpaired t-test: ChR2 vs YFP t(4) = 3.25, p = 0.053)] (Fig. 5D-F). In contrast, c-Fos+ cell density was significantly increased in DBH-cells in ChR2 rats compared to YFP [whole VLM (n = 4-5/group, unpaired t-test: ChR2 vs YFP t(7) = 3.67, p = 0.016)] (Fig. 5C). This effect was present in the rostral, intermediate, and caudal subregions of the VLM [rostral VLM (unpaired t-test: ChR2 vs YFP t(5) = 5.49, p = 0.0055); intermediate VLM (unpaired t-test: ChR2 vs YFP t(6) = 3.04, p = 0.048); caudal VLM (unpaired t-test: ChR2 vs YFP t(5) = 4.016, p = 0.023)] (Fig. 5D-F).

Similar trends were observed in females. ChR2 stimulation increased the density of c-Fos+ DBH-cells in the VLM as a whole, as well as within the rostral and intermediate subregions [whole VLM (n = 4-6/group, unpaired t-test: ChR2 vs YFP t(8) = 3.34, p = 0.022); rostral VLM (unpaired t-test: ChR2 vs YFP t(8) = 4.49, p = 0.0049); intermediate VLM (unpaired t-test: ChR2 vs YFP t(6) = 3.23, p = 0.036) (Fig. 5G-I). Although, there was no change in cFos+ DBH-cell density in the caudal VLM (unpaired t-test: ChR2 vs YFP t(3) = 2.50, p = 0.17)] (Fig. 5J). Similar to males, there were no effects on c-Fos+ cell density of DBH+ neurons in any female VLM region [whole VLM (n = 4-6/group, unpaired t-test: ChR2 vs YFP t(7) = 1.58, p = 0.32); rostral VLM (unpaired t-test: ChR2 vs YFP t(4) = 1.63, p = 0.16); intermediate VLM (unpaired t-test: ChR2 vs YFP t(3) = 1.79, p = 0.16); caudal VLM (unpaired t-test: ChR2 vs YFP t(6) = 0.21, p > 0.99)] (Fig. 5G-J). Collectively, these data demonstrate that vmPFC terminal stimulation preferentially activates non-catecholaminergic neurons throughout the VLM in both sexes.

### vmPFC Inputs Target Medullary Inhibitory Neurons

Next, we sought to identify the non-catecholaminergic VLM neurons targeted by the vmPFC. Prior work has demonstrated that a local network of inhibitory neurons acts on VLM catecholaminergic neurons to regulate sympathoexcitation (Gao et al., 2019; Guyenet et al., 1990; Heesch et al., 2006). In fact, GABAergic and glycinergic neurons are recruited by barosensitive-neurons in the nucleus of the solitary tract to regulate RVLM outflow and control blood pressure (Guyenet, 2006; Schreihofer & Guyenet, 2002). Here, GABA and GlyT2, a marker for glycinergic neurons, were immunolabeled and YFP-expressing vmPFC fibers were found to appose GABAergic and glycinergic neurons in the RVLM (Fig. 7A). Additionally, optogenetic stimulation led to c-Fos expression in GABAergic RVLM neurons (Fig. 7B). The current data in aggregate support a model where vmPFC projections to inhibitory RVLM neurons provide a mechanism for limiting catecholaminergic output and subsequent physiological stress responses (Fig. 7C).

**Figure 7.**
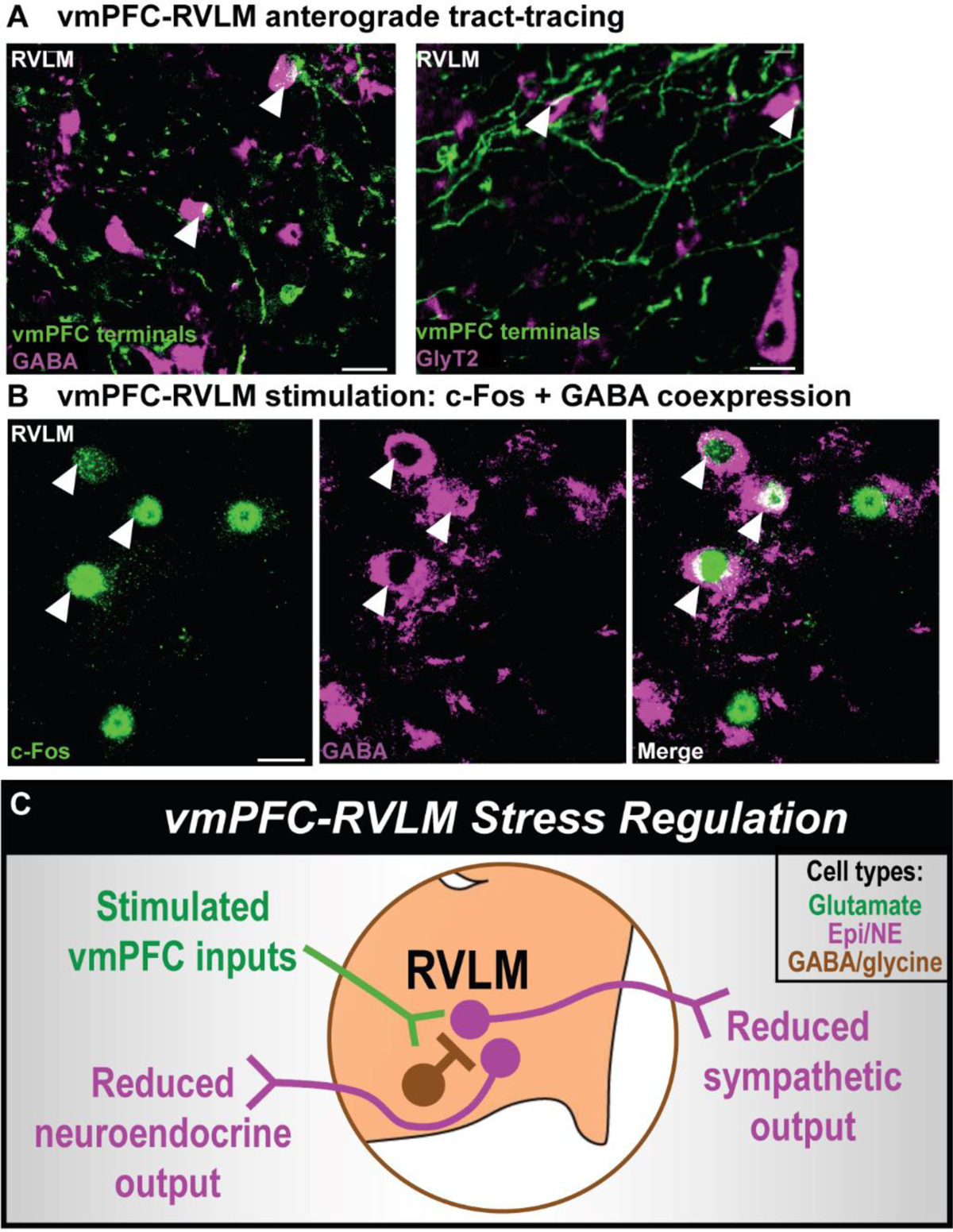
vmPFC-RVLM circuit mechanism. vmPFC terminals expressing YFP apposed GABA- and GlyT2-expressing RVLM cells, arrowheads denote appositions, scale bars: 20 μm (A). ChR2 rats were immunolabeled for c-Fos and GABA following vmPFC-RVLM stimulation, GABA+ and c-Fos+ cells are marked by arrowheads, scale bars: 20 μm (B). A schema illustrating a proposed mechanism for the vmPFC-RVLM circuit (C). Epi/NE: Epinephrine/Norepinephrine-synthesizing neurons.

## Discussion

The current studies mapped the structural and functional connectivity of a prefrontal-medullary circuit and identified a role in stress response inhibition. Here, vmPFC terminals apposed catecholaminergic neurons throughout the VLM. Additionally, stress-reactive vmPFC neurons projected to the RVLM and CVLM through a mixture of parallel and divergent pathways. vmPFC-RVLM circuit activation also reduced glucocorticoid stress reactivity in both male and female rats, while stress-induced hyperglycemia was blunted only in males.

However, the vmPFC-RVLM circuit did not mediate preference or aversion behavior in either sex. The vmPFC-RVLM circuit preferentially activated non-catecholaminergic neurons and targeted GABAergic and glycinergic neurons in the RVLM. Collectively, these studies reveal a vmPFC-RVLM circuit that inhibits endocrine stress responses in male and female rats, potentially via RVLM inhibitory neurons.

### vmPFC Regulation of Stress Responding

Human neuroimaging studies have identified the vmPFC as an integral hub for emotion, cognition, goal-directed behavior, and physiological regulation (Jennings et al., 2016; Kraynak et al., 2018; Nestler et al., 2002; Saper & Stornetta, 2014). Additionally, vmPFC gray matter and functional activity are frequently altered in depression and other stress-related disorders that disproportionately burden females (Drevets et al., 2008; Riecher-Rössler, 2017). Accordingly, rodent studies have examined the vmPFC to identify neural substrates contributing to sex-specific vulnerability (Hurley & Carelli, 2020; van der Zee et al., 2022; Woodward et al., 2023). In vivo monitoring of vmPFC projection neurons also revealed that sex and stress history interact to shape neural activity in response to both appetitive and aversive stimuli (Wallace & Myers, 2023). Further, stimulation of vmPFC glutamate neurons produces opposing outcomes in male and female rats. Male vmPFC neuronal stimulation reduces HPA axis and autonomic responses to stress, while also enhancing sociability and motivation (Wallace et al., 2021). Female vmPFC stimulation facilitates stress responses, including hyperglycemia and tachycardia, but does not affect social motivation (Wallace et al., 2021). Mechanisms for cortical regulation of stress responding may lie in downstream circuitry. The current report found that the vmPFC-RVLM pathway inhibits physiological stress responding in both sexes without effects on preference behavior. Therefore, this projection to the brainstem may account, in part, for the stress-inhibitory effects of male vmPFC (Wallace et al., 2021). However, the current results do not align with prior reports of female cortical stress excitation (Wallace et al., 2021). In all, the vmPFC-RVLM circuit inhibits stress responding in both sexes, a largely sex-similar function.

### Importance of Biological Sex

Although our findings demonstrate vmPFC-RVLM activation reduces stress reactivity in both sexes, differences may still exist between sexes. Male and female data were not statistically compared due to experiments occurring at different times. However, there appear to be magnitude differences in circuit structure and function as anterograde/retrograde mapping and c-Fos expression point toward more robust connectivity in males. It is possible that the sex-specific glycemic regulation reported relates to differences in circuit strength. However, it is difficult to disentangle potential effects of biological sex from necessary differences in experimental parameters. There were subtle differences in the volume of viral constructs administered to generate similar anatomical coverage in the targeted nuclei of the male and female brain. Hence, although the general properties of circuit organization and function were similar in males and females, multiple factors prevent direct comparison of male and female results and limit interpretation of sex-specific outcomes such as glucose mobilization. Nonetheless, there are reports of sex differences in sympathoexcitation and endogenous glucose production is higher in males than females (Amiel et al., 1993). Sex differences in sympathetic activity likely involve the RVLM as estrogen receptors alpha (ERα) and beta (ERβ) are present in catecholaminergic and non-catecholaminergic RVLM neurons of female rats (Hay, 2016; Saleh & Connell, 2000; Wang et al., 2006). Interestingly, ultrastructural imaging found immunoreactive-ERβ predominately in extra-nuclear sites, while ERα localizes to the nucleus (Wang et al., 2006). Additionally, 17β-estradiol acting through ERβ inhibits voltage-gated calcium currents in bulbospinal RVLM neurons and RVLM ERβ knockdown in female mice exacerbates aldosterone/salt-induced hypertension (Xue et al., 2013). ERβ signaling also elicits transient vasodepressor effects in male rats (Shih, 2009). Together, these studies demonstrate that ERβ limits neurogenic sympathetic outflow, possibly through a nongenomic mechanism in RVLM neurons. Other hormone receptors including androgen receptors are present in RVLM catecholaminergic neurons and glia of male and female rodents (Milner et al., 2007; Sheng et al., 2021), but little is known about the impact on stress physiology.

### Regulation of RVLM Catecholaminergic Neurons

Catecholaminergic neurons in the RVLM are often defined as C1 neurons and characterized by the presence of phenylethanolamine N-methyl transferase (PNMT), the synthesis enzyme for epinephrine. These epinephrine-synthesizing neurons target preganglionic sympathetic neurons in the spinal cord to trigger the sympathomedullary (SAM) axis and induce physiological responses such as elevating arterial pressure and blood glucose (Guyenet, 2006; Guyenet et al., 2013). Additionally, caudal RVLM and CVLM catecholaminergic neurons are often labeled as A1 neurons and characterized by the presence of DBH and the absence of PNMT (Stornetta, 2009). These norepinephrine-synthesizing neurons target the paraventricular hypothalamus to influence HPA axis output (Guyenet et al., 2013; Ritter, 2017). When stimulated, medullary catecholaminergic neurons increase SAM and HPA axes activity prompting corticosterone and glucose synthesis and release (Li et al., 2018; Zhao et al., 2017). In this study, glucose and corticosterone were measured to evaluate SAM and HPA axes function during stress. To restrain physiological stress responses, outputs from medullary catecholaminergic neurons are tonically inhibited by GABAergic and glycinergic VLM neurons (Gao et al., 2019; Guyenet et al., 1990; Heesch et al., 2006). This microcircuitry is engaged by the baroreflex to regulate sympathetic output and maintain blood pressure homeostasis (Guyenet, 2006), providing an intrinsic network for cortical modulation.

Prior ultrastructural studies identified vmPFC synapses in the RVLM (Gabbott et al., 2007), yet the functional organization of this circuitry has not been reported. In this study, stimulating the vmPFC-RVLM circuit increased c-Fos expression preferentially in non-catecholaminergic neurons. Although we cannot rule out activation of non-catecholaminergic glutamatergic neurons, this is a relatively small cell population with the majority of RVLM catecholaminergic neurons expressing vGluT2 (Stornetta et al., 2002; Stornetta, 2009). Thus, the identified vmPFC inputs to VLM inhibitory populations represent a potential mechanism for cortical inhibition of stress responding. Nevertheless, the widespread vmPFC innervation of multiple VLM cell-types suggests the circuit is dynamically regulated and that different physiological or environmental states may lead to divergent circuit signaling and homeostatic regulation. Ultimately, the potential for differential vmPFC inputs to catecholaminergic and non-catecholaminergic VLM cells may provide a cellular basis for stimulus-specific physiological reactivity.

### The vmPFC-RVLM in Brain-Body Function

Responding to challenges requires the coordination of autonomic and neuroendocrine systems for optimal physiological adaptation. Considering the current results, the vmPFC-RVLM appears to be an effective means for modulating physiological function. Conventionally, physiological response patterns have been proposed to involve networks of top-down neural pathways that integrate cortical and limbic information to provide inputs to midbrain nuclei that ultimately modulate descending brainstem outflow. Yet, a monosynaptic cortical-medullary pathway provides a rapid and direct pathway for controlling homeostatic functions. As the vmPFC is essential for contextual appraisal and emotional processing, while also processing visceral information (Kraynak et al., 2018; Saper, 2002; Verberne & Owens, 1998), this circuit may provide a direct link for psychosomatic coordination that relates emotion and physical health.

## Conclusion

The current studies advance understanding of how cognitive appraisal of stressors impacts physiological function. Specifically, vmPFC inputs to the RVLM restrain glucocorticoid stress reactivity, likely through the activation of non-catecholaminergic RVLM neurons. Additional investigation of cortical glutamate signaling within the microcircuitry of the medulla is likely to identify novel aspects of endocrine-autonomic integration that could represent therapeutic interventions to improve cardiometabolic health.

## Additional Information Section

### Author Contributions

Pace SA and Myers B designed the research plan; Pace SA, Lukinic E, Wallace T, McCartney C, and Myers B. performed the experiments; Pace SA and Wallace T analyzed the data; Pace SA and Myers B wrote the manuscript, and all authors approved the final submitted version.

## Acknowledgments

AAV5-packaged vectors for ChR2 and YFP were provided by the University of North Carolina Vector Core under material transfer agreement with Karl Deisseroth and Stanford University. AAVretro vectors expressing mCherry were a gift from Karl Deisseroth (Addgene viral prep # 114472-AAVrg) and the GFP-expressing AAVretro was a gift from Bryan Roth (Addgene viral prep # 50465-AAVrg). The authors thank Jacob Moore, Derek Schaeuble, and Carley Dearing for technical support, and Courtney Bouchet for manuscript editing. This article was first published as a preprint: Pace SA, Lukinic E, Wallace T, McCartney C, Myers B. Cortical-brainstem circuitry attenuates stress reactivity. bioRrxiv

## References

Amiel, S. A., Maran, A., Powrie, J. K., Umpleby, A. M., & Macdonald, I. A. (1993). Gender differences in counterregulation to hypoglycaemia. Diabetologia, 36(5), 460–464. 10.1007/BF00402284

Bekhbat, M., Glasper, E. R., Rowson, S. A., Kelly, S. D., & Neigh, G. N. (2018). Measuring corticosterone concentrations over a physiological dynamic range in female rats. Physiology and Behavior, 194, 73–76. 10.1016/j.physbeh.2018.04.033

Bialik, R. J., Smythe, J. M., & Roberts, D. C. S. (1988). Alpha2-adrenergic receptors mediate the increase in blood glucose levels induced by epinephrine and brief footshock stress. Progress in Neuropsychopharmacology and Biological Psychiatry, 12(2–3), 307–314. 10.1016/0278-5846(88)90049-8

Card, J. P., Sved, J. C., Craig, B., Raizada, M., Vazquez, J., & Sved, A. F. (2006). Efferent projections of rat rostroventrolateral medulla C1 catecholamine neurons: Implications for the central control of cardiovascular regulation. Journal of Comparative Neurology, 499(5), 840–859. 10.1002/cne.21140

Cora, M. C., Kooistra, L., & Travlos, G. (2015). Vaginal Cytology of the Laboratory Rat and Mouse:Review and Criteria for the Staging of the Estrous Cycle Using Stained Vaginal Smears. Toxicologic Pathology, 43(6), 776–793. 10.1177/0192623315570339

Dearing, C., Morano, R., Ptaskiewicz, E., Mahbod, P., Scheimann, J. R., Franco-Villanueva, A., Wulsin, L., & Myers, B. (2021). Glucoregulation and coping behavior after chronic stress in rats: Sex differences across the lifespan. Hormones and Behavior, 136, 105060. 10.1016/j.yhbeh.2021.105060

Drevets, W. C., Price, J. L., & Furey, M. L. (2008). Brain structural and functional abnormalities in mood disorders: implications for neurocircuitry models of depression. Brain Structure and Function, 213(1–2), 93–118. 10.1007/s00429-008-0189-x

Drevets, W. C., Price, J. L., Simpson, J. R., Todd, R. D., Reich, T., Vannier, M., & Raichle, M. E. (1997). Subgenual prefrontal cortex abnormalities in mood disorders. Nature, 386(6627), 824–827. 10.1038/386824a0

Duncan, J. (2001). An adaptive coding model of neural function in prefrontal cortex. Nature Reviews Neuroscience, 2(11), 820–829. 10.1038/35097575

Fuchikami, M., Thomas, A., Liu, R., Wohleb, E. S., Land, B. B., DiLeone, R. J., Aghajanian, G. K., & Duman, R. S. (2015). Optogenetic stimulation of infralimbic PFC reproduces ketamine’s rapid and sustained antidepressant actions. Proceedings of the National Academy of Sciences of the United States of America, 112(26), 8106–8111. 10.1073/pnas.1414728112

Gabbott, P. L. A., Warner, T. A., Jays, P. R. L., Salway, P., & Busby, S. J. (2005). Prefrontal cortex in the rat: Projections to subcortical autonomic, motor, and limbic centers. Journal of Comparative Neurology, 492(2), 145–177. 10.1002/cne.20738

Gabbott, P. L. A., Warner, T. A., & Busby, S. J. (2007). Catecholaminergic neurons in medullary nuclei are among the post-synaptic targets of descending projections from infralimbic area 25 of the rat medial prefrontal cortex. Neuroscience. 144(2), 623–635. 10.1016/j.neuroscience.2006.09.048

Gao, H., Korim, W. S., Yao, S. T., Heesch, C. M., & Derbenev, A. v. (2019). Glycinergic neurotransmission in the rostral ventrolateral medulla controls the time course of baroreflex-mediated sympathoinhibition. Journal of Physiology, 597(1), 283–301. 10.1113/JP276467

Guyenet, P. G. (2006). The sympathetic control of blood pressure. In Nature Reviews Neuroscience (Vol. 7, Issue 5, pp. 335–346). Nature Publishing Group. 10.1038/nrn1902

Guyenet, P. G., Darnall, R. A., & Riley, T. A. (1990). Rostral ventrolateral medulla and sympathorespiratory integration in rats. American Journal of Physiology - Regulatory Integrative and Comparative Physiology, 259(5 28–5). 10.1152/ajpregu.1990.259.5.r1063

Guyenet, P. G., Stornetta, R. L., Bochorishvili, G., DePuy, S. D., Burke, P. G. R., & Abbott, S. B. G. (2013). C1 neurons: The body’s EMTs. In American Journal of Physiology - Regulatory Integrative and Comparative Physiology (Vol. 305, Issue 3). 10.1152/ajpregu.00054.2013

Hamani, C., Mayberg, H., Stone, S., Laxton, A., Haber, S., & Lozano, A. M. (2011). The subcallosal cingulate gyrus in the context of major depression. In Biological Psychiatry (Vol. 69, Issue 4, pp. 301–308). Elsevier. 10.1016/j.biopsych.2010.09.034

Hay, M. (2016). Sex, the brain and hypertension: Brain oestrogen receptors and high blood pressure risk factors. In Clinical Science (Vol. 130, Issue 1, pp. 9–18). Portland Press. 10.1042/CS20150654

Heesch, C. M., Laiprasert, J. D., & Kvochina, L. (2006). RVLM glycine receptors mediate GABAA and GABAB independent sympathoinhibition from CVLM in rats. Brain Research, 1125(1), 46–59. 10.1016/j.brainres.2006.09.090

Hurley, K. M., Herbert, H., Moga, M. M., & Saper, C. B. (1991). Efferent projections of the infralimbic cortex of the rat. Journal of Comparative Neurology, 308(2), 249–276. 10.1002/cne.903080210

Hurley, S. W., & Carelli, R. M. (2020). Activation of infralimbic to nucleus accumbens shell pathway suppresses conditioned aversion in male but not female rats. Journal of Neuroscience, 40(36), 6888–6895. 10.1523/JNEUROSCI.0137-20.2020

Jennings, J. R., Sheu, L. K., Kuan, D. C. H., Manuck, S. B., & Gianaros, P. J. (2016). Resting state connectivity of the medial prefrontal cortex covaries with individual differences in high-frequency heart rate variability. Psychophysiology, 53(4), 444–454. 10.1111/psyp.12586

Kraynak, T. E., Marsland, A. L., & Gianaros, P. J. (2018). Neural Mechanisms Linking Emotion with Cardiovascular Disease. In Current Cardiology Reports (Vol. 20, Issue 12, pp. 1–10). Current Medicine Group LLC 1. 10.1007/s11886-018-1071-y

Li, A. J., Wang, Q., & Ritter, S. (2018). Selective pharmacogenetic activation of catecholamine subgroups in the ventrolateral medulla elicits key glucoregulatory responses. Endocrinology, 159(1), 341–355. 10.1210/en.2017-00630

Liotti, M., Mayberg, H. S., Brannan, S. K., McGinnis, S., Jerabek, P., & Fox, P. T. (2000). Differential limbic– cortical correlates of sadness and anxiety in healthy subjects: implications for affective disorders. Biological Psychiatry, 48(1), 30–42. 10.1016/S0006-3223(00)00874-X

McKlveen, J. M., Myers, B., & Herman, J. P. (2015). The Medial Prefrontal Cortex: Coordinator of Autonomic, Neuroendocrine and Behavioural Responses to Stress. In Journal of Neuroendocrinology (Vol. 27, Issue 6, pp. 446–456). 10.1111/jne.12272

Milner, T. A., Hernandez, F. J., Herrick, S. P., Pierce, J. P., Iadecola, C., & Drake, C. T. (2007). Cellular and subcellular localization of androgen receptor immunoreactivity relative to C1 adrenergic neurons in the rostral ventrolateral medulla of male and female rats. Synapse, 61(5), 268–278. 10.1002/syn.20370

Myers, B., McKlveen, J. M., & Herman, J. P. (2014). Glucocorticoid actions on synapses, circuits, and behavior: Implications for the energetics of stress. In Frontiers in Neuroendocrinology (Vol. 35, Issue 2, pp. 180–196). 10.1016/j.yfrne.2013.12.003

Myers, B., McKlveen, J. M., Morano, R., Ulrich-Lai, Y. M., Solomon, M. B., Wilson, S. P., & Herman, J. P. (2017). Vesicular glutamate transporter 1 knockdown in infralimbic prefrontal cortex augments neuroendocrine responses to chronic stress in male rats. Endocrinology, 158(10), 3579–3591. 10.1210/en.2017-00426

Nestler, E. J., Barrot, M., DiLeone, R. J., Eisch, A. J., Gold, S. J., & Monteggia, L. M. (2002). Neurobiology of depression. In Neuron (Vol. 34, Issue 1, pp. 13–25). Cell Press. 10.1016/S0896-6273(02)00653-0

Pace, S. A., Christensen, C., Schackmuth, M. K., Wallace, T., McKlveen, J. M., Beischel, W., Morano, R., Scheimann, J. R., Wilson, S. P., Herman, J. P., & Myers, B. (2020). Infralimbic cortical glutamate output is necessary for the neural and behavioral consequences of chronic stress. Neurobiology of Stress, 13, 100274. 10.1016/j.ynstr.2020.100274

Paxinos, G., & Watson, C. (2006). The Rat Brain in Stereotaxic Coordinates. 10.1016/0143-4179(83)90049-5

Riecher-Rössler, A. (2017). Sex and gender differences in mental disorders. In The Lancet Psychiatry (Vol. 4, Issue 1, pp. 8–9). Elsevier. 10.1016/S2215-0366(16)30348-0

Ritter, S. (2017). Monitoring and Maintenance of Brain Glucose Supply. In Appetite and Food Intake (pp. 177– 204). CRC Press/Taylor & Francis. 10.1201/9781315120171-9

Ritter, S., Li, A. J., & Wang, Q. (2019). Hindbrain glucoregulatory mechanisms: Critical role of catecholamine neurons in the ventrolateral medulla. In Physiology and Behavior (Vol. 208, p. 112568). NIH Public Access. 10.1016/j.physbeh.2019.112568

Saleh, T. M., & Connell, B. J. (2000). 17beta-estradiol modulates baroreflex sensitivity and autonomic tone of female rats. Journal of Autonomic Nervous System, 80(3), 148–161. 10.1016/S0165-1838(00)00087-4

Saper, C. B. (2002). The central autonomic nervous system: Conscious visceral perception and autonomic pattern generation. In Annual Review of Neuroscience (Vol. 25, pp. 433–469). Annual Reviews. 10.1146/annurev.neuro.25.032502.111311

Saper, C. B., & Stornetta, R. L. (2014). Central Autonomic System. In The Rat Nervous System: Fourth Edition (pp. 629–673). Elsevier. 10.1016/B978-0-12-374245-2.00023-1

Schaeuble, D., Packard, A. E. B., McKlveen, J. M., Morano, R., Fourman, S., Smith, B. L., Scheimann, J. R., Packard, B. A., Wilson, S. P., James, J., Hui, D. Y., Ulrich-Lai, Y. M., Herman, J. P., & Myers, B. (2019). Prefrontal Cortex Regulates Chronic Stress-Induced Cardiovascular Susceptibility. Journal of the American Heart Association, 8(24), e014451. 10.1161/JAHA.119.014451

Schreihofer, A. M., & Guyenet, P. G. (2002). The baroreflex and beyond: Control of sympathetic vasomotor tone by gabaergic neurons in the ventrolateral medulla. Clinical and Experimental Pharmacology and Physiology, 29(5–6), 514–521. 10.1046/j.1440-1681.2002.03665.x

Sheng, J. A., Tan, S. M. L., Hale, T. M., & Handa, R. J. (2021). Androgens and Their Role in Regulating Sex Differences in the Hypothalamic/Pituitary/Adrenal Axis Stress Response and Stress-Related Behaviors. Androgens: Clinical Research and Therapeutics, 2(1), 261–274. 10.1089/ANDRO.2021.0021

Shih, C. D. (2009). Activation of estrogen receptor - dependent nitric oxide signaling mediates the hypotensive effects of estrogen in the rostral ventrolateral medulla of anesthetized rats. Journal of Biomedical Science, 16(1), 60. 10.1186/1423-0127-16-60

Solomon, M. B., Karom, M. C., & Huhman, K. L. (2007). Sex and estrous cycle differences in the display of conditioned defeat in Syrian hamsters. Hormones and Behavior, 52(2), 211–219. 10.1016/j.yhbeh.2007.04.007

Stamatakis, A. M., & Stuber, G. D. (2012). Activation of lateral habenula inputs to the ventral midbrain promotes behavioral avoidance. Nature Neuroscience, 15(8), 1105–1107. 10.1038/nn.3145

Stornetta, R. L. (2009). Neurochemistry of bulbospinal presympathetic neurons of the medulla oblongata. In Journal of Chemical Neuroanatomy (Vol. 38, Issue 3, pp. 222–230). NIH Public Access. 10.1016/j.jchemneu.2009.07.005

Stornetta, R. L., & Guyenet, P. G. (2018). C1 neurons: a nodal point for stress? Experimental Physiology, 103(3), 332–336. 10.1113/EP086435

Stornetta, R. L., Inglis, M. A., Viar, K. E., & Guyenet, P. G. (2016). Afferent and efferent connections of C1 cells with spinal cord or hypothalamic projections in mice. Brain Structure and Function, 221(8), 4027–4044. 10.1007/s00429-015-1143-3

Swanson, L. W. (2004). Brain maps: structure of the rat brain (Issue 3rd edition). 10.1016/0166-2236(93)90187-q

Ulrich-Lai, Y. M., & Herman, J. P. (2009). Neural regulation of endocrine and autonomic stress responses. Nature Reviews: Neuroscience, 10, 397–409.

van der Zee, Y. Y., Lardner, C. K., Parise, E. M., Mews, P., Ramakrishnan, A., Patel, V., Teague, C. D., Salery, M., Walker, D. M., Browne, C. J., Labonté, B., Parise, L. F., Kronman, H., Penã, C. J., Torres-Berrío, A., Duffy, J. E., de Nijs, L., Eijssen, L. M. T., Shen, L., … Nestler, E. J. (2022). Sex-Specific Role for SLIT1 in Regulating Stress Susceptibility. Biological Psychiatry, 91(1), 81–91. 10.1016/j.biopsych.2021.01.019

Verberne, A. J. M., & Owens, N. C. (1998). Cortical Modulation of theCardiovascular System. Progress in Neurobiology, 54(2), 149–168. 10.1016/S0301-0082(97)00056-7

Wallace, T., & Myers, B. (2023). Prefrontal representation of affective stimuli: importance of stress, sex, and context. Cerebral Cortex. 10.1093/CERCOR/BHAD110

Wallace, T., Schaeuble, D., Pace, S. A., Schackmuth, M. K., Hentges, S. T., Chicco, A. J., & Myers, B. (2021). Sexually divergent cortical control of affective-autonomic integration. Psychoneuroendocrinology, 129, 2020.09.29.319210. 10.1016/j.psyneuen.2021.105238

Wang, G., Drake, C. T., Rozenblit, M., Zhou, P., Alves, S. E., Herrick, S. P., Hayashi, S., Warrier, S., Iadecola, C., & Milner, T. A. (2006). Evidence that estrogen directly and indirectly modulates C1 adrenergic bulbospinal neurons in the rostral ventrolateral medulla. Brain Research, 1094(1), 163–178. 10.1016/j.brainres.2006.03.089

Wood, M., Adil, O., Wallace, T., Fourman, S., Wilson, S. P., Herman, J. P., & Myers, B. (2019). Infralimbic prefrontal cortex structural and functional connectivity with the limbic forebrain: a combined viral genetic and optogenetic analysis. Brain Structure and Function, 224(1), 73–97. 10.1007/s00429-018-1762-6

Woodward, E., Coutellier, L., Rangel-Barajas, C., Ringland, A., & Logrip, M. L. (2023). Sex-Specific Timelines for Adaptations of Prefrontal Parvalbumin Neurons in Response to Stress and Changes in Anxiety-and Depressive-Like Behaviors. ENeuro, 10(3). 10.1523/ENEURO.0300-22.2023

Xue, B., Zhang, Z., Beltz, T. G., Johnson, R. F., Guo, F., Hay, M., & Johnson, A. K. (2013). Estrogen receptor-β in the paraventricular nucleus and rostroventrolateral medulla plays an essential protective role in aldosterone/salt-induced hypertension in female rats. Hypertension, 61(6), 1255–1262. 10.1161/HYPERTENSIONAHA.111.00903

Zhao, Z., Wang, L., Gao, W., Hu, F., Zhang, J., Ren, Y., Lin, R., Feng, Q., Cheng, M., Ju, D., Chi, Q., Wang, D., Song, S., Luo, M., & Zhan, C. (2017). A Central Catecholaminergic Circuit Controls Blood Glucose Levels during Stress. Neuron, 95(1), 138–152.e5. 10.1016/j.neuron.2017.05.031

